# When is a slender not a slender? The cell-cycle arrest and scarcity of slender parasites challenges the role of bloodstream trypanosomes in infection maintenance

**DOI:** 10.1101/2023.04.06.535835

**Authors:** Stephen D. Larcombe, Emma M. Briggs, Nick Savill, Balazs Szoor, Keith R. Matthews

## Abstract

The development of *Trypanosoma brucei* in its mammalian host is marked by a distinct morphological change as replicative “slender” forms differentiate into cell-cycle arrested “stumpy” forms in a quorum-sensing dependent manner. Although stumpy forms dominate chronic infections at the population level, the proportion of replicative parasites at the individual cell level and the irreversibility of arrest in the bloodstream is unclear. Here, we experimentally demonstrate that developmental cell cycle arrest is definitively irreversible in acute and chronic infections in mice. Furthermore, analysis of replicative capacity and single-cell transcriptome profiling reveals a temporal hierarchy, whereby cell-cycle arrest and appearance of a stumpy-like transcriptome precedes irreversible commitment and morphological change. Unexpectedly, we show that proliferating parasites are exceptionally scarce in the blood after infections are established. This challenges the ability of bloodstream trypanosomes to sustain infection by proliferation or antigenic variation, these parasites instead being overwhelmingly adapted for transmission.

## Introduction

African trypanosomes are protozoan parasites that cause sleeping sickness in humans and ‘nagana’ in livestock, responsible for a significant humanitarian and economic burden in sub Saharan Africa(Giordani et al., 2016). These species mostly require a tsetse fly vector to complete their life cycles, necessitating a suite of complex developmental changes(Rotureau B and Van Den Abbeele, 2013). Atypically among the trypanosome group, the development of *Trypanosoma brucei* in its mammalian hosts is accompanied by a distinct morphological change; replicative ‘slender’ forms establish the infection, before differentiating into non-dividing ‘stumpy’ forms through quorum sensing(Bruce et al., 1912). These stumpy forms are cell cycle arrested, more resistant to destruction by the host immune system(McLintock et al., 1993), and exhibit molecular pre-adaptation for their onward development in tsetse flies(Matthews, 2021). This morphological transition has been noted since the earliest descriptions of these parasites(Bruce et al., 1912; Robertson, 1912), and much recent work has focussed on uncovering the molecular signalling events that underpin the developmental changes(Rojas and Matthews, 2019). Nonetheless, fundamental questions about the commitment to differentiation of bloodstream forms in the mammalian host and the contribution of different developmental forms to the infection dynamic remain.

In particular, a trade-off is thought to exist between the parasites’ investment in slender and stumpy forms. Specifically, the proliferation of slender forms maintains the infection and promotes immune evasion by enabling the generation and expansion of antigenic variants in the parasite population(Mugnier, 2015). In contrast, the generation of stumpy forms limits the parasitaemia and so promotes host survival, whereas their molecular adaptations prepare them for their transmission to tsetse(Matthews and Larcombe, 2021). Being non-proliferative and uniformly arrested in their cell cycle in G1/G0, stumpy forms of *T. bruce*i have been considered terminally arrested and destined to be eliminated unless taken up by a tsetse fly, where they can resume proliferation as differentiating procyclic forms(Szoor et al., 2020). This combination of immune evasion by antigenic variation, the proliferation of slender forms and the accumulation and then senescence or immune elimination of arrested stumpy forms has long been associated with a classic profile of undulating waves of trypanosome parasitaemias. However, despite long-standing acceptance that, once initiated, the transition from slender to stumpy form is terminal, evidence for this is indirect (Cunningham et al., 1963; Dewar et al., 2018; Hamm et al., 1990; Lumsden, 1972; Reuner et al., 1997; Turner et al., 1995; Vassella et al., 1997). Moreover, trypanosome parastiaemias are pleomorphic, reflecting that a progressive spectrum of morphologies exists between slender and stumpy extremes, this being supported by recent single cell transcriptomic analyses (Briggs et al., 2021). Hence the relationship between cell proliferation, morphological and molecular adaption, and the reversibility of cell cycle arrest for bloodstream parasites is unclear. Added to this complexity, most experimental studies of slender and stumpy forms have focused only on the first wave of infection whereas chronic infections are more representative of clinical trypanosomiasis in humans and animals (Macgregor et al., 2011).

Here, using an ex vivo assay of parasite proliferation using wild type and quorum-sensing defective parasites, and single cell transcriptomic analysis, we have quantitatively analysed the proportion and molecular characteristics of replicable or arrested parasites at acute and chronic stages of infection, as well as parasites transitional between the morphological extremes. Surprisingly, our data suggest that, regardless of morphology, the bloodstream parasite population has an insignificant role in the maintenance of the infection, instead being devoted to transmission.

## Results

### Bloodstream trypanosomes lose replicative competence soon after infection

To explore the commitment of trypanosomes to developmental arrest in the bloodstream we used an ex vivo plating assay, whereby pleiomorphic *T. brucei* AnTat 90:13 parasites from infections were carefully counted, serially diluted, and then the proportion of parasites able to proliferate determined by their outgrowth in vitro. This established their ‘replicative competence’, i.e. the percentage of proliferating cells, or cells able to resume proliferation in vitro, at different stages of an infection in mice over 24 days (Figure 1a). Figures 1a and 1b demonstrate that early in the establishment phase of the infection (<120h post infection) there was an excellent correlation between the number of parasites in vivo and their replicative competence (100%±0%). This confirmed the ability of the assay to accurately quantitate replication competent cells in the infection and demonstrated that the parasites were uncommitted to arrest at this point. Between 120h and 128h post-infection, however, the proportion of replication competent parasites declined rapidly to 13.7±11.1% by 144h, this coinciding with the peak of parasitaemia in the first wave of infection (Figure 1a). Following this, between 192 and 288h the parasites recrudesced and 27.5±6.6% were replication competent. Beyond this point (384h-576h), however, the parasitaemia fluctuated between 1×10^7^ and 10^8^ but the proportion of replication competent cells never exceeded 5.0±4.3% (Figure 1a).

**Figure 1.**
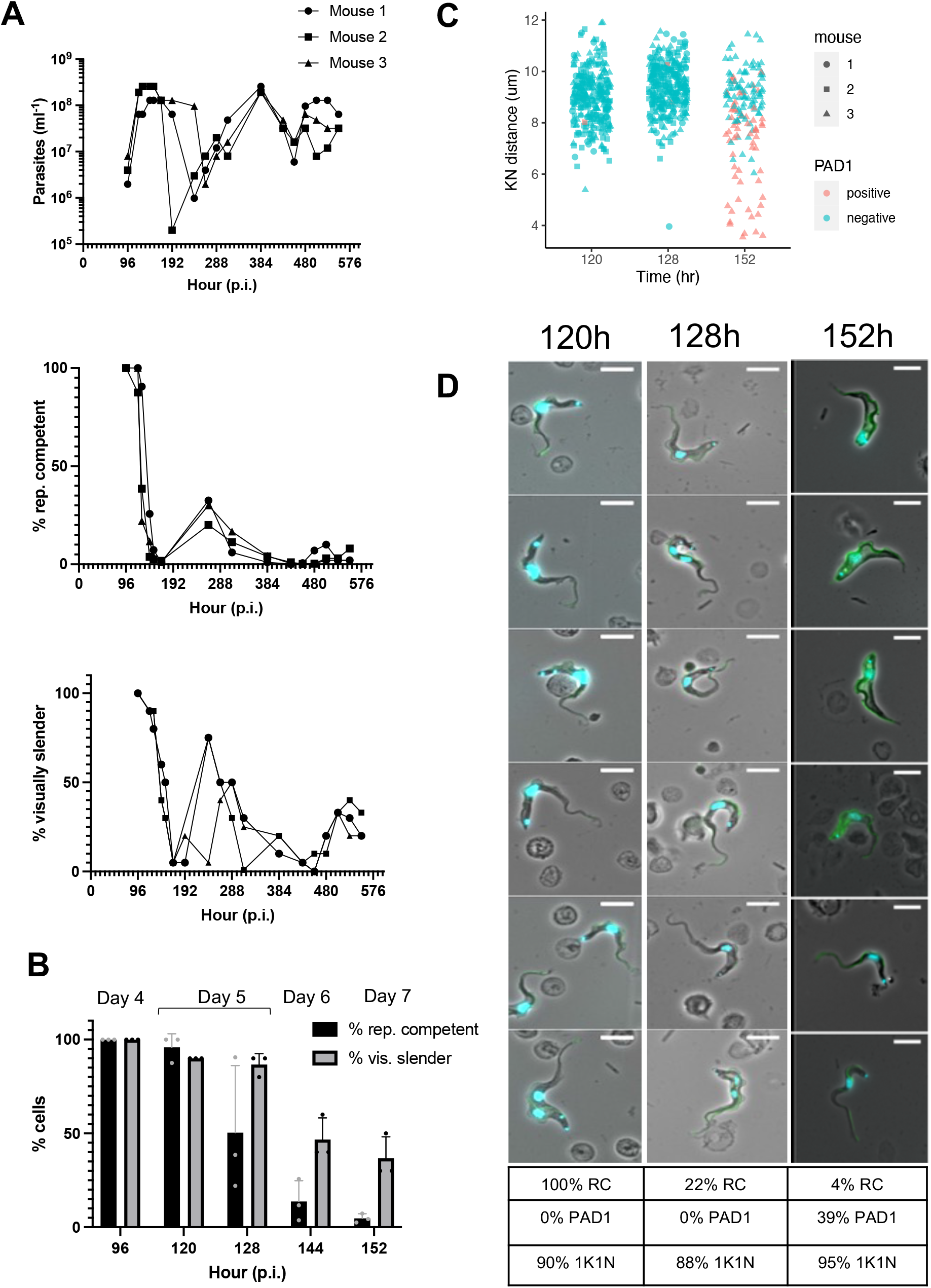
Non-proliferative forms dominate mouse infections. a) Parasitaemia, % replication (rep.) competent cells and morphology at different timepoints from a wild type *T. brucei* Antat 1.1 90:13 infection of mice (n=3) b) Comparison of the frequency of replication competent cells and those with slender morphology post infection (p.i.) (n=3) c) Analysis of the kinetoplast-nucleus dimension at 120h, 128h and 152h post infection; PAD+ cells are in pink. d) Representative phase contrast images of parasites at 120h,128h and 152h of infection, counterstained for nucleus and kinetoplast (blue) and PAD1 (green). All cells are morphologically slender and PAD1 -ve at 120h and 128h. PAD1+ve stumpy and PAD1-slender cells are present at 152h. The Table shows the % replication competent (RC), PAD1 +ve and cells with 1 kinetoplast and 1 nucleus (1K1N) for the respective infections. Scale bar; 10μm.

Replicative competence describes the proportion of parasites able to proliferate, whether derived from replicative, morphologically slender cells or from non-replicative, morphologically slender or stumpy forms, that are not irreversibly committed to arrest and which can resume growth in vitro. Therefore, at each time point during infection we related the morphology of the parasites to their replicative competence to assess the relationship between cell morphology and the commitment to arrest. Figure 1b demonstrates that until 120h the population was overwhelmingly slender in morphology, matching their replicative competence. However, between 128 and 152h morphologically slender cells always exceeded cells with replicative competence, demonstrating that the parasites had committed to arrest prior to their morphological transformation. These changes were very rapid, such that parasites in one mouse dropped from being 100% to 22% replication competent between 120-128h post infection (Fig 1b, d). Beyond 152h, when stumpy cells were prevalent, the proportion of morphologically slender cells varied between 50% and 1%. Nonetheless, even when up to 30% of visually slender cells were detected late in the chronic phase of infection, only 2.5% of parasites exhibited replicative competence (Figure 1a). For a more detailed assessment of morphology and developmental change, we investigated the expression of the stumpy marker protein PAD1(Dean et al., 2009), and the kinetoplast to nucleus (KN) distance, focussing on the 120h and 128h timepoints when the rapid change in replicative competence occurred. Figures 1c and 1d show that there was no difference in either measure, or overall cell morphology, at these timepoints, despite the fall in replicative competence, contrasting with 152h, when there was a clear change in both PAD1 expression and KN distance. Further images of fields of cells at 120h, 128h and 456h are available in Supplementary Figure 1.

In combination, these assays demonstrated, firstly, that the commitment to replicative arrest precedes PAD1 expression and morphological development to stumpy forms, secondly, that stumpy forms cannot resume proliferation once removed from the density-sensing signal in the blood and, thirdly, that, after the early phase of the infection, replication competent parasites in the bloodstream are scarce as a proportion of the population.

### Disrupting the quorum sensing pathway increases replicative competence

An alternative explanation for the loss of replicative competence during infections was a failure to grow in media after adaptation to the host environment. To eliminate this possibility, we exploited a genetic perturbation that reduces the quorum sensing capacity of the parasites and so delays the development from slender to stumpy forms. We predicted this would increase replicative competence of parasites later in infection if adaptation to the host was not an important factor for their growth ex vivo. Therefore, we initiated infections with a parasite line engineered for doxycycline regulated RNAi-mediated silencing of HYP2, an identified component of the trypanosome quorum sensing signalling pathway (Mony et al., 2014; Silvester et al., 2017). In the first peak (Figure 2a left) non-induced cells behaved as wild type controls, with a reduction in replicative competence preceding the morphological change of the parasites. With HYP2 RNAi silencing, in contrast, the loss of replicative competence was reduced (F=5.77, p=0.003) as was the morphological transition (F=6.5, p=0.007), consistent with reduced quorum sensing mediated differentiation (Figure 2a, b).

**Figure 2.**
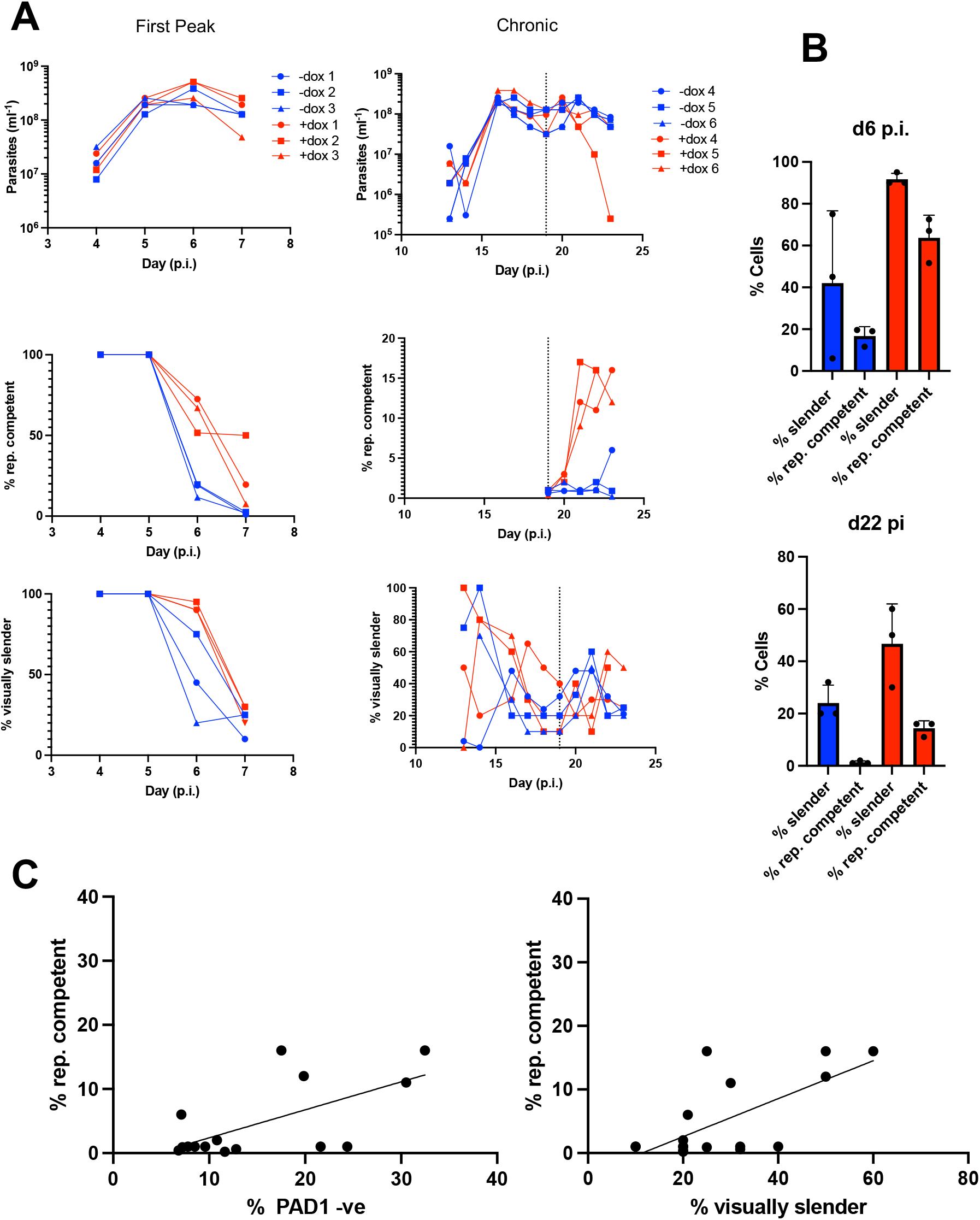
RNAi against HYP2 delays commitment to stumpy formation. a) Parasitaemia, morphology and % replication competent cells in the first peak (left) or chronic phase (right) of infection with (+dox; red) or without (-dox; blue) induction of RNAi against HYP2. The dashed line indicates the point that induction was started with doxycycline b) Percentage of cells with characteristics plotted in a, isolated after d6 p.i. or d22 p.i. to highlight show the treatment differences between -dox (blue) and + dox (red). c) Comparison between the proportion of replication competent and PAD1 -ve (left; p<0.05) or visually slender (right; p=0.09) cells.

The consequences of HYP2 RNAi were further evaluated in the chronic phase of the infections, between d19 and d23, when replicative competence is very low. (Figure 2a, right). Although there was variation between mice, by d22 post infection there was an increase in both morphologically slender cells and those exhibiting replicative competence (non-induced 1.33%±0.57; induced 14.33%±2.88; F_4,18_=12.81, p<0.0001; Figure 2a right, Figure 2b). Extending the analysis beyond morphology for both induced and uninduced parasites further demonstrated that PAD1 expression (F=15.48, p=0.002) and slender morphology (F=3.31, p=0.09) showed an inverse correlation with replicative competence but that the relationship was not 1:1; many PAD1 negative, morphologically slender cells, were not replication competent (Figure 2c).

Thus, more slender form parasites derived from acute and chronic infections can successfully proliferate in vitro if quorum sensing is reduced, eliminating host adaptation as an explanation for the low replicative capacity in wild type cells. Instead, regardless of their morphology, after initial establishment parasites show extremely limited replicative competence due to their commitment to development to stumpy forms.

### The stumpy-associated gene expression programme is activated before irreversible commitment

To investigate the characteristics of cell types (replicative slender, arrested stumpy and uncommitted or committed cells of any morphology) in acute and chronic stages of infection, we exploited single cell RNA sequencing with the Chromium platform (Briggs et al., 2021). Parasites were isolated from day 7 and day 23 of infection, with HYP2 RNAi being induced or not 5 days prior to harvesting (i.e. at day 2 in acute infection and day 19 for the chronic infections). After quality control filtering 3826 transcriptomes remained for d7-dox, 7721 for d7+dox, 8435 for d23-dox and 7721 for d23+dox. All samples were integrated prior to dimension reduction and clustering analysis (Supplementary Table 1).

Across all four samples we identified 5 clusters of cells defined with distinct marker genes (Figure 3a). To broadly characterise the clusters, we used expression levels of a panel of existing marker genes (Briggs et al., 2021) to assign “slender” and “stumpy” scores to each cell, and also the respective clusters were labelled according to cell cycle marker gene expression (Figure 3b, Supplementary Figure 3). As expected from the numbers of replication competent cells at both d7 and d23, only a small minority of parasites showed high expression of slender marker genes which nearly exclusively belonged to cluster 3 (Figure 3c). This cluster, which we consider “true” slender cells comprised < 2.7% of cells in any sample and was the only cluster exhibiting expression of markers consistent with active cell cycle progression (Figure 3b; Supplementary Figure 3). All the other clusters showed low slender scores and higher expression of stumpy marker genes, although cluster 4 and cluster 1 parasites showed notably lower stumpy scores than cluster 2 and cluster 0. Cluster 0, the most common group of cells in all samples, is associated with high stumpy scores, and represents terminally committed stumpy forms. Consistent with this, these cells were marked by relatively high expression, for example, of EP and GPEET procyclins, ZC3H20, PAD2 and ZFPs, each of which has been found to be upregulated as parasites prepare for transmission or undergo stumpy formation(McWilliam et al., 2019; Szoor et al., 2020). The cyclin F box2 protein was also elevated, this being associated with VSG mRNA stability(Bravo Ruiz et al., 2022).

**Figure 3.**
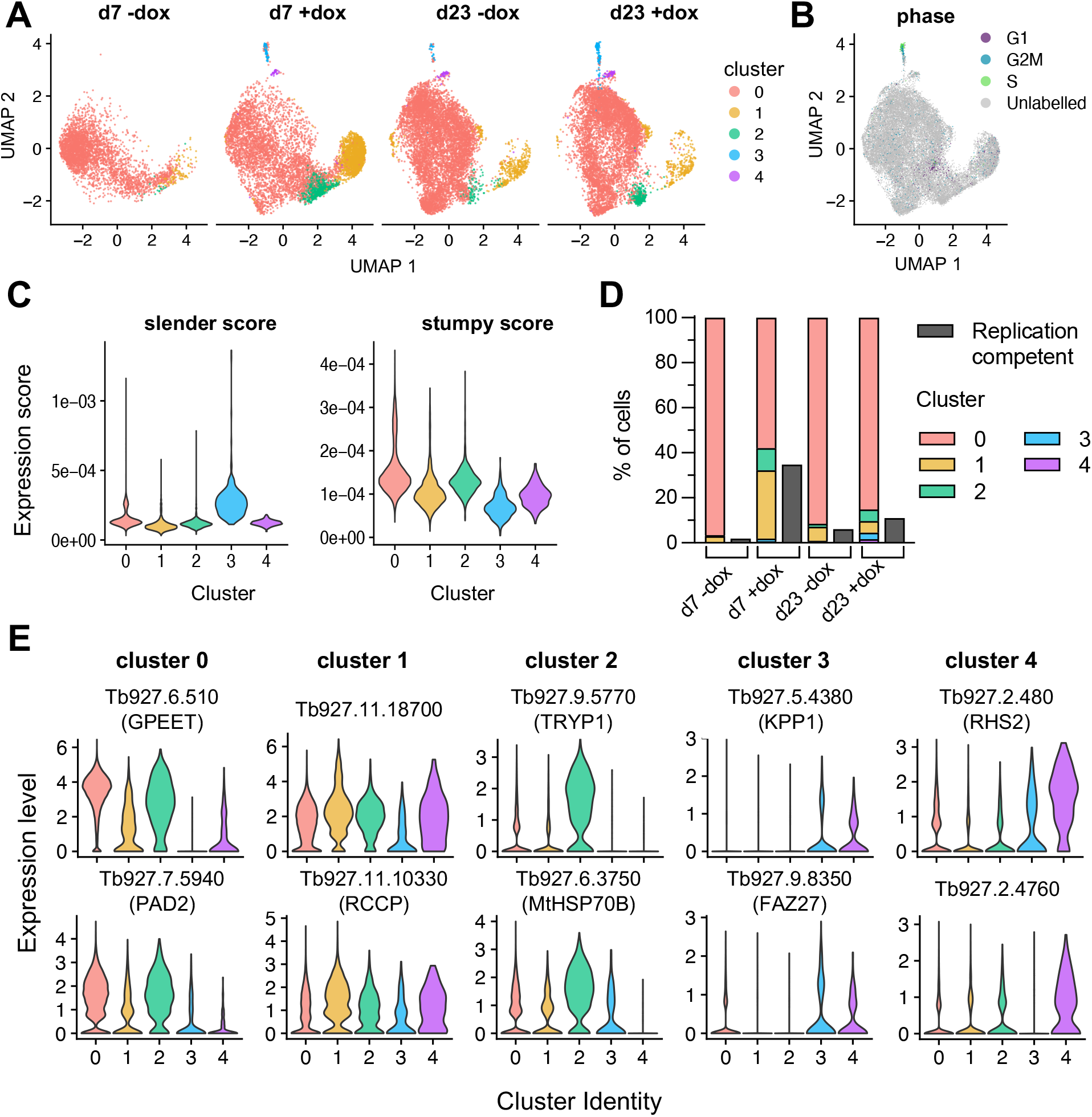
Single cell RNA-seq reveals the scarcity of parasites with a slender-associated transcriptome. a) UMAP plots of cell transcriptomes at day 7 and day 23 post-infection, ± tet induction of HYP2 RNAi. Each point is one cell, coloured by cluster identity. Key to the right is applicable to each plot. b) Cells labelled by cycle phase, for all samples c) The “slender score” and “stumpy score” consisting of the average expression of associated marker genes for each cluster identified in panel a, for all samples. d) Comparison of the proportion of each cluster with the % of replication competent cells in each sample e) Expression of two cluster enriched marker genes for clusters 0-5, analysed with respect to each cluster.

Importantly, the proportion of cells in each sample that was replication competent after ex vivo plating was always higher than the proportion of true slender cells (cluster 3), implicating the presence of cells in other clusters that were not yet irreversibly committed to arrest (Figure 3d). By examining the cumulative representation of distinct clusters in each experimental condition (acute, chronic, HYP2 induced or not), we found that a composite score, comprised of a combination of cells belonging to clusters 1, 2 and 4, provided an excellent match to the corresponding proportion of cells that were replication competent in each sample (Figure 3d). Thus, their number increased with HYP2 RNAi and was lower when HYP2 was not silenced, supporting their identification as uncommitted cells. Critically, despite the anticipation that they are replication competent, the transcriptomic profiles of cells belonging to clusters 1 and 4 do not match those of cluster 3, the true slender cells, suggesting they have embarked on the developmental programme to stumpy forms. Interestingly, although cluster 1 is similar to the transcriptome profile of stumpy forms (cluster 0), it exhibited higher expression of Tb927.11.18700, a reported target of the Grumpy snoRNA implicated in stumpy development (Guegan et al., 2022), a chromatin remodelling ISWI complex component (Stanne et al., 2011) and a telomere associated protein (Reis et al., 2018) linked to expression site silencing, a feature of development to stumpy forms(Amiguet-Vercher et al., 2004). Notably, no cell cycle markers were elevated in clusters 1, 2 and 4 (Figure 3b; Supplementary Figure 3) demonstrating an absence of their replicative activity in the bloodstream, despite the ability of these uncommitted cells to resume growth when isolated and cultured ex vivo. Comparative datasets for each cluster with respect to the overall population are available for interrogation in Supplementary Table 1, whereas enriched marker transcripts for each cluster are shown in Figure 3e.

In combination, these data quantitatively reveal that there is an exceptionally small population of true slender cells in active replication in the blood, together with a population of uncommitted cells that do not proliferate, share the expression of many genes expressed in stumpy forms and yet are capable of re-entering the cell cycle if removed from the bloodstream environment (Figure 4). Finally, the remaining and dominant proportion in the bloodstream, are terminally arrested stumpy forms.

**Figure 4.**
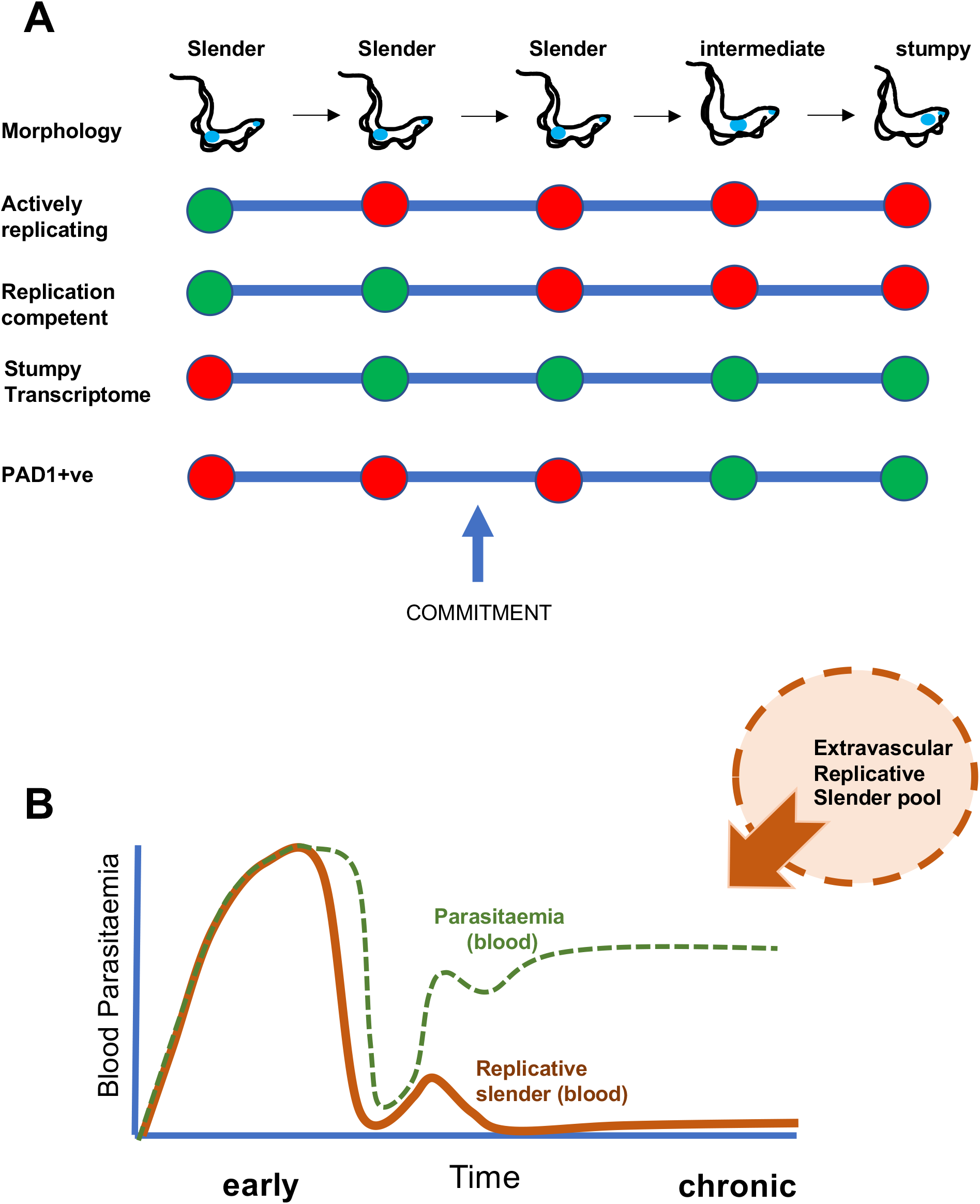
The developmental commitment of slender forms and their contribution to the infection dynamic. a) Events in the development of slender forms toward stumpy forms, highlighting that the acquisition of a stumpy-like transcriptome and irreversible commitment precedes morphological change and PAD1 protein expression. B) Model for the source of blood parasites given the scarcity of replication competent slender forms in the vasculature.

## Discussion

The established view of trypanosomes in the mammalian bloodstream is that slender forms proliferate to sustain the infection and generate antigenic diversity whilst stumpy forms are irreversibly arrested and preadapted for transmission for tsetse flies. The direct evidence for both of these elements of the infection dynamic however is limited. In this study, we found that morphologically stumpy forms were incapable of re-entering proliferation if removed from the quorum sensing signal. Moreover, by careful temporal analysis during the first wave of parasitaemia, we could establish that the commitment to arrest precedes morphological transformation such that cells indistinguishable from proliferative bloodstream slender forms early in the parasitaemia lost replicative competence significantly before morphologically stumpy forms appeared. By day 7, single cell analysis showed not one of 3826 cells (d7-dox) had a transcriptome representative of true slender (cluster 3) cells (Briggs et al., 2021), and cells with replicative competence were generally present at less than 2% of the population. Analysis in the chronic phase of infection confirmed that stumpy forms adapted for transmission dominated, as previously proposed based on population level analysis (Macgregor et al., 2011), but importantly revealed at the level of individual cells that slender forms capable of proliferation were exceptionally rare, representing only 0.33% of the bloodstream trypanosome population at d22. Moreover, by single cell RNA sequence analysis, only around two thirds of even this small proportion appeared to be undergoing active proliferation. Overall, we can conclude that slender cells undergo irreversible commitment to stumpy formation before any morphological change (Figure 4a) and that the proliferation of slender forms in the bloodstream makes a much smaller than anticipated contribution to the maintenance of infection (Figure 4b). Indeed, the proportion of replicative slender forms is even less at the individual cell level than previously estimated based on PAD1 expression and mathematical modelling (Macgregor et al., 2011).

Previous analyses of the irreversibility of stumpy cell cycle arrest are based on mathematical modelling of the parasitaemia (Macgregor et al., 2011; Seed and Black, 1997) and the inference that stumpy enriched populations are quantitatively less infective to naïve animals (Cunningham et al., 1963). In our assays, the ability to proliferate ex vivo was analysed and the parasites were found to precipitously lose replicative competence from approximately 120h of infection in mice, whereafter it remained very low. We considered that this could be contributed to by the parasites becoming adapted to the mammalian host environment and so less able to efficiently proliferate ex vivo. This possibility, however, was eliminated by silencing a component of the quorum sensing pathway; under these conditions, the capacity of the cells to become stumpy was reduced in vivo and, correspondingly, the replicative competence of the parasites increased. Importantly this was the case not only early in infection but also 20 days after growth of the parasites in vivo. Thus, the parasites develop to stumpy forms during the course of the parasitaemia and once generated, these cells cannot re-enter proliferation and are terminally committed to cell cycle arrest, at least until development to the next life cycle stage in the tsetse fly. Definitively, therefore, stumpy formation is irreversible.

From quantitative analysis of replication competence and the transcriptome profile of the parasite population, replicative slender cells frequently represent less than 1% of trypanosomes in the chronic phase of infection, which is more typical of natural long term infections. Is this sufficient for bloodstream parasites to sustain the infection? Although many parameters are uncertain, our data suggests not. For example, between d19-d21 the proportion of replication competent cells never exceeded 1%, which could not generate the two-fold increase in total parasitaemia observed during this chronic period, particularly when combined with an expected decay in the stumpy forms through senescence (Dewar et al., 2018). One explanation to sustain the infection is that the population is replenished from niches elsewhere. Characterised extravascular locations for trypanosomes include the skin, lungs and adipose tissue(Capewell et al., 2016; Mabille et al., 2022; Trindade et al., 2016), and the migration between the adipose tissues and blood has recently been estimated to be 11% of blood parasites/per milligram of tissue/day, highlighting that there is significant, though restricted, potential for exchange(Trindade et al., 2022). These extravascular compartments have recently been identified as sites of elevated antigenic diversity (Beaver et al., 2023) and, if this is combined with enhanced replicative capacity, could seed parasites into the bloodstream where they rapidly arrest and transition to stumpy forms. This creates a model where the parasites disseminated in the tissues serve to populate the circulation with a pool of parasites optimised to transmit to haematophagous tsetse flies.

Single cell RNA-sequencing allowed the molecular characteristics of parasites in the bloodstream population to be interrogated in the early and chronic phase of infection for the first time. Moreover, inclusion in the analysis of cells after HYP2 silencing informed on the relationship between the transcriptome profile of the respective cell clusters and their replicative potential. Specifically, beyond the majority of cells with a stumpy transcriptome (cluster 0) and the tiny proportion of true slender cells (cluster 3), there was a significant group of cells (cluster 1, 2) that were similar to, but distinct from, stumpy forms plus a small group of cells similar to replicative slender forms (Cluster 4). Consistently, the combined proportion of these clusters was equivalent to the proportion of cells which retained proliferative competence ex vivo (with their proportion increasing as quorum sensing is reduced by HYP2 silencing). We interpret these clusters as representing cells which are not committed to irreversible arrest and yet have undergone significant developmental adaptation. Although able to proliferate ex vivo if removed from the quorum sensing signal, these parasites would remain arrested in vivo and progress to full commitment and stumpy formation without regaining replicative capacity. This generates a hierarchy of development whereby morphologically slender forms are initially proliferative, then reversibly activate a transcriptome profile similar to stumpy forms, followed by irreversible commitment to arrest and finally morphological transition to the stumpy form (Figure 4a).

Our data addresses several key aspects of the infection biology of trypanosomes in the mammalian blood. Firstly, it formally demonstrates that the development of trypanosomes to stumpy forms entails terminal developmental arrest in the mammalian host. Secondly, by temporal analysis of replicative competence through the first wave of parasitaemia and single cell transcriptome analysis, the commitment to arrest and development was found to precede PAD1 expression and morphological change to stumpy forms. Thirdly, our discovery that replicative slender forms in the bloodstream are exceptionally rare challenges the ability of the bloodstream parasite population to maintain the infection or drive antigenic variation, complementing the description of elevated antigenic diversity in extravascular niches (Beaver et al., 2023). Furthermore, although it has been reported that slender, PAD1-negative, cells have the potential for onward development in flies (Schuster et al., 2021), our data suggests many such cells would in fact be already committed to stumpy development regardless of their morphology and any replicative slender cells would be insufficiently prevalent to contribute meaningfully to transmission. Finally, by the single cell analysis of parasite populations early and late in the infection we provide a platform for the biological analysis of parasites in the chronic stage of infection. These have been almost entirely neglected to date but are most relevant for infections in the field.

## Author contributions

KRM: acquired funding, designed the study, analysed the data, wrote the manuscript SL: designed the study, carried out parasite experiments, analysed the data, wrote the manuscript

EB: designed the study, carried out scRNA experiments and data processing, analysed the data, wrote the manuscript

BS: carried out parasite experiments, analysed the data

NS: Interpreted the data and performed quantitative analysis.

## Materials availability

All unique/stable reagents generated in this study are available from the Lead Contact without restriction.

## Data and code availability

Original code is available at. https://tinyurl.com/28ydamyy

For scRNAseq data, the raw fastq are registered with the ENA (https://www.ebi.ac.uk/ena/browser/home) under the study reference - PRJEB60851

Then authors declare no competing interests

## Methods

### Mouse Infections

In vivo experiments were performed under UK Home Office licence number (PP2251183) approved after review at the University of Edinburgh ethical review committee.

For all experiments we used 10-12 week old female Balb/c mice (Charles River). For Wild Type infections, mice were infected with 100 cells of a pleiomorphic *T*.*brucei* EATRO1125 AnTat 1.1 90:13 (Engstler and Boshart, 2004). For the HYP2 RNAi infections mice were infected with 100 cells of a previously described *T. brucei* EATRO1125 AnTat1.1 90:13 cell line with a doxycycline-inducible RNAi targeting *Tb*HYP2(Mony et al., 2014). In both cases frozen stocks were cultured for 2 days prior to infection to ensure all cells in the inoculum were viable. Three experiments were carried out using the inducible HYP2 RNAi line: in the first experiment induction of RNAi commenced on d0 of infection and was terminated at d7 p.i.. In the second RNAi against HYP2 was induced on d19 of infection and was terminated on d23. In the third experiment, RNAi against HYP2 was induced on d19 of infection and terminated on d28. In all cases induction was achieved by addition of doxycycline to drinking water (200μg/ml in 5% sucrose). Non-induced controls received drinking water with 5% sucrose. In all mouse infections, mice were blood sampled regularly following d3 p.i. by tail snip. Parasitaemia was assessed using Herbert and Lumsden matching charts for fields of view at 40x magnification (Herbert and Lumsden, 1976). Morphology was also scored: the number of clearly slender parasites was counted for at least three fields of view and used to create a % of slender cells for each day. Slender assignment was scored based on the morphology, the length of flagella and the motility of cells, and was always done within 30 minutes of blood sampling. All blood sampling was kept within UK Home Office limits as specified in the relevant project licence. In addition to rapid scoring on days for analysis of replication competence we also kept around 5ul of blood in 500ul of HMI-9 media. This was kept warm for subsequent use in plating assays and for making blood smears.

### Plating assay

We developed a plating assay for assessment of *ex vivo* proliferation of parasites from mice. Following blood sampling and parasitaemia scoring, we used the diluted blood sample in media to fill plates with known numbers of cells. We used an improved Neubauer haemocytometer to accurately count the parasites in each 500μl. The counts were used to create a dilution for a final stock with a fixed concentration of 1 cell/μl in 5ml total volume. Since parasitaemia is commonly around 1 × 10^7^ cells per ml in blood, this rapid dilution to 1000 per ml represents a 10,000 fold dilution, effectively removing the density dependent oligopeptide differentiation signal. The stock was then used to fill a round bottom 96 well plate with the following cell numbers: 50/well, 20/well, 5/well, 2/well and 1/well. The wells were filled with HMI-9 media to 100ul total volume. In the HYP2 RNAi experiment we continued doxycycline treatment from mice into the 96 well plate. After seeding the plates were left untouched for 5-7 days then each well was assessed for an outgrowing culture of parasites in binary fashion. After 5-7 days there was almost always nothing, or a viable culture in each well. On rare occasions where only a few cells were in a well, it was left a further three days and then re-scored. Wells around the edges of the plate were not used as pilot work indicated these grew less successfully. Thus, there were 12 replicates for each seeding. The number of wells were counted for each seeding that developed outgrowth, and the percentage growing for each condition was calculated. Since only one competent cell is required for outgrowth, probability (P) theory was used to calculate the estimated proportion of replication competent cells using the following formula: P(at least one cell replicative) = 1 -P(failure in one trial)^n^ where n is the total number of trials. We used an average of the values of replication competent cells across the seedings where there was variation in outgrowth, avoiding those where everything or nothing grew: these seeding do not offer enough variation to create an accurate estimate.

### Microscopy

In addition to rapid scoring of “slenderness” for a subset of samples leftover cells from the plating assay were used to examine the types of cell present in the samples in detail with immunofluorescent microscopy. These were 120 hours and 128 hours post infection in the WT experiment, and days 5, 7, 19, 22 and 23 in our HYP2 RNAi experiment. On each of these days, after quenching any mouse antibodies for at least 30 mins in HMI-9, the blood was centrifuged (10 mins at 400g) and the supernatant removed. The remaining blood and parasite pellet was used to make 2 blood smears per mouse. Smears were air dried before being fixed and stored in methanol at -20°C. Cell cycle analysis and PAD1 protein expression analysis were carried out by staining ice-cold methanol fixed cells with 4’,6-diamidino-2-phenylindole (DAPI) (100 ng/ ml) and an anti-PAD1 antibody (Dean et al., 2009) as previously described (Silvester et al., 2017). Images were made of all slides at 63x magnification, aiming for at least 100 cells for each slide. The cell cycle status was determined by counting the number of kinetoplasts and nuclei in each cell (KN score) and scoring for presence/absence of staining with the PAD1 antibody. Positive cells were classed based on any PAD1 staining that was not restricted to only the flagella or flagellar pocket. For a non-binary description of differentiation, the kinetoplast to nucleus distance (KN distance) was also measured from all images of cells in 1K1N this providing a quantitative proxy for the change from slender to stumpy as the cell reconfigures morphologically. Using imageJ (Schindelin et al., 2012) a script was used to automatically measure the distance between the centres of two particles in the DAPI stained images. The distance in pixels was converted to μm by dividing by 6.106155.

### Statistical Analyses

For the mouse experiments, three animals per treatment were used. With effects sizes similar to those previously observed for RNAi mediated loss of developmental competence (0.637 to 1.804; e.g.(Mony et al., 2014)) a sample size of three per treatment provided sufficient power to test for treatment differences. To test for differences in replicative competence mediated by treatment a one-way repeated measures ANOVA was constructed with interest in the time* treatment (+ or – dox) interaction. To assess how differences in characteristics at the population level (%2K1N, 2K2N*P; % PAD1 +ve*; % visually slender) were related to the percentage of replication competent cells for a given timepoint, a multiple linear regression analysis was constructed. This model assessed the significance of relationships between each variable, while holding treatment groups as fixed factors. The models were simplified by removing non-significant terms until only interactions with a p value of < 0.1 remained. In all cases the simplified models provided a better fit to the observed data than larger models, as assessed by sum of squares (smaller is better). Blinding was not carried out. P values of less than 0.05 were considered statistically significant in all cases. For scRNA-seq differential expression tests, a significance threshold of adjusted p-value < 0.05 was used.

### Chromium (10x Genomics) library preparation and Illumina sequencing

Parasites were purified from terminal bleed sample using diethylaminoethyl-cellulose columns(Lanham and Godfrey, 1970). Following isolation from host cells into PSG buffer (1X PBS + 1% D-glucose, pH 7.8), centrifugation at 400 x g for 10 min was used to pellet parasites which were resuspended in 1 ml HMI-9 supplemented with 20% FCS. Parasites were counted with haemocytometer and adjusted to ∼ 2×10^6^ cells/ml. Parasites were then mixed 1:1 with 2X freezing media (HMI-9 with 20% glycerol) and frozen at -80°C wrapped in cotton wool to slow freezing. After 24-72 hr, parasites were moved to liquid nitrogen storage. For Chromium scRNA-seq (10x Genomics), sample were transported on dry ice before slow defrosting to limit cell death using previously validated protocol(Emma M. Briggs et al., 2023). For each sample, parasites were combined from 2-3 mice in equal proportion depending on sample viability, only samples with > 90% live, motile, parasites were used for sequencing. Samples day 7 -dox (n=3) and +dox (n=2) were processed in one batch. The first run of day 23 -dox (n=3) and day 23 +dox (n=2) samples resulted in low cell recovery; 1084 and 623 cells, respectively. Therefore, a second batch of remaining samples (n=1 for -dox and n =2 for +dox) were processed for day 23, and two batches were combined for day 23 samples. Sample preparation was performed with the 3’ Gene Expression kit (version 3.1 chemistry) and sequencing was performed with Illumina NextSeq 2000, to generate 28 bp x 130 bp paired reads to high depth: 62,083 means reads per cell for day 7 -dox, 61,345 mean reads per cell for day 7 +dox, 81,559 mean reads per cell for day 23 minus -dox and 91,441 mean reads per cell for day 23 +dox. All fastq files are available on ENA under project PRJEB60851.

### scRNA-seq data processing and analysis

Data was mapped to the *T. brucei* TREU927 with extended UTR sequences (Briggs et al., 2021) using Cellranger software version 7 to generate counts matrices. As the day 7 -dox sample showed higher than expect number of droplets with low total RNA, we controlled for potential free RNA in the samples with SoupX (Young and Behjati, 2020). Resulting counts were further quality control filtered to remove low quality transcriptomes and likely multiplets (Supplementary Figure 2). For each sample, variable genes were selected before data was log2 normalised and scaled using previously described methods (Briggs et al 2021). All four samples were integrated using Seurat version 4 (Hao et al., 2021) before dimension reduction and clustering analysis was performed. Differential expression testing between clusters was performed using MAST (Finak et al., 2015). Average expression scores were calculated using the AddModuleScore function from Seurat which finds the average expression levels of marker gene set across each cell, subtracted by the aggregated expression of a randomly selected control gene set (Tirosh et al., 2016). For cell cycle phase scores, previously published marker genes were used, and early and late G1 markers were combined (Archer et al., 2011). To generate slender and stumpy marker gene sets, the single cell transcriptomes of *in vitro* generated slender and stumpy forms (Briggs et al., 2021) were compared to find genes with highest fold change (> 2) with MAST differential expression test, resulting in 414 slender and 110 stumpy marker genes. All code and counts matrices are available on Zenodo (10.5281/zenodo.7778583).

## Supplementary Figures and Tables

**Supplementary Figure 1**

Representative images of fields of parasites from 120h, 128h and 456h post infection counterstained with DAPI (light blue) and PAD1 green. Individual parasites are highlighted in red boxes.

**Supplementary Figure 2**

Metadata for the single cell RNA seq analysis of each sample (d7 +dox; d7 -dox; d23 +DOX; d23 -dox) before and after quality control filtering.

**Supplementary Figure 3**

scRNA-seq reveals the majority of blood form *T. brucei* lack expression of cell cycle phase marker genes. a) Average expression score of G1 (left), S (centre) and G2/M (right) phase marker genes for each parasite from all samples (D7 -dox and +dox, and D23 -dox and +dox). Parasites are split by clusters (0-4, x-axis). Red dashed line show threshold applied (0.05), to assign a phase to each parasite. Cells with expression below 0.05 in all case where marked “Unlabelled”. b) UMAPs showing parasites for each samples (left to right), coloured by the assigned cell cycle phase. c) The percentage of cells in each cluster (x-axis) assigned in each cell cycle phase. d) The percentage of cells from each sample (x-axis), in each cell cycle phase (left bars), as well as the percentage of cells able to replicate when removed from host (i.e. replication competent cells) (dark grey, right bars).

**Supplementary Table 1**

Cluster marker genes identified by differential expression analysis for each cluster. Significant marker genes (p_val_adj < 0.05) over expressed in one cluster compared to all others are listed.

