## supplementary figures for "When is a slender not a slender? The cell-cycle arrest and scarcity of slender parasites challenges the role of bloodstream trypanosomes in infection maintenance"

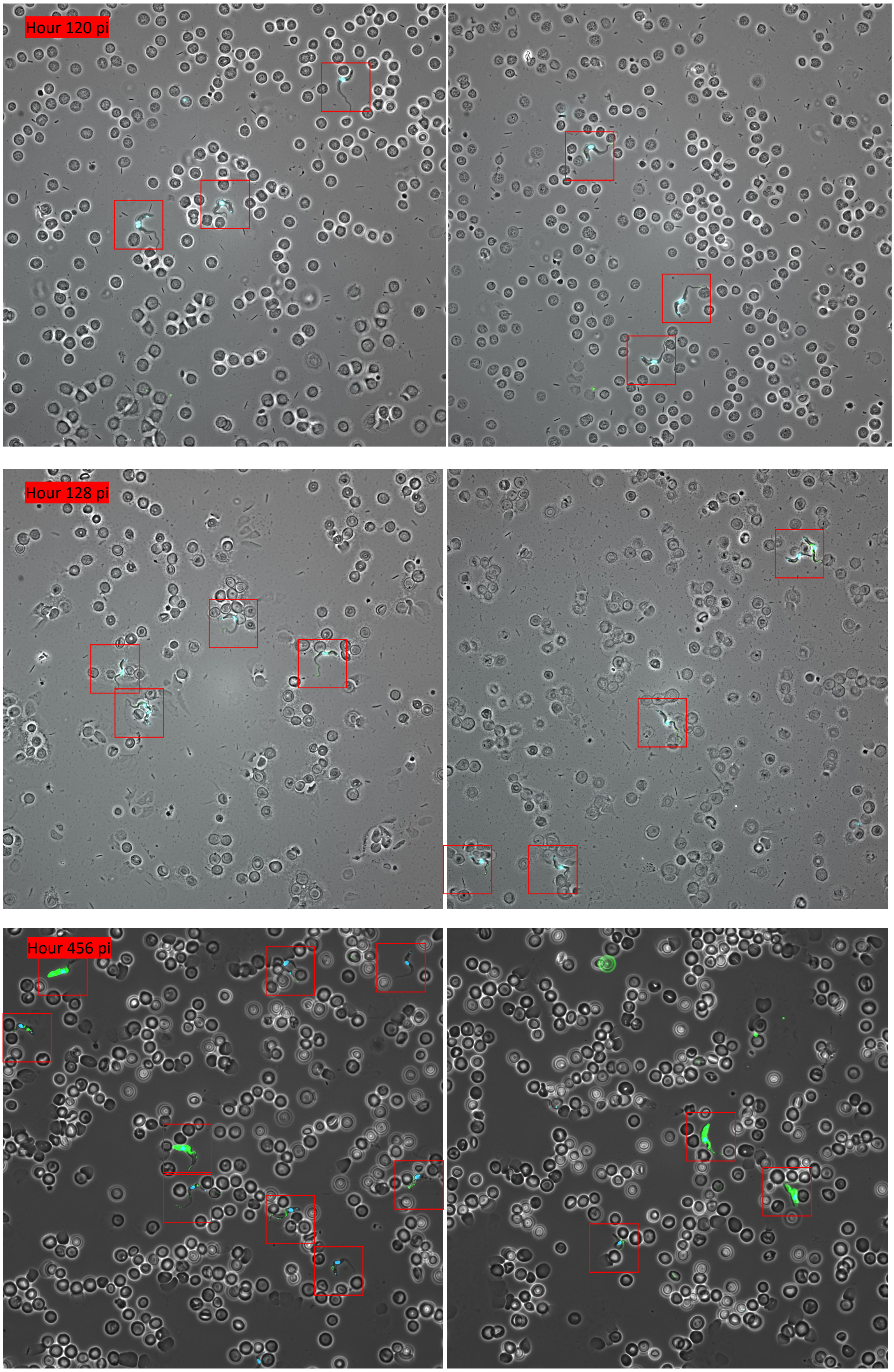


**Supplementary Figure 1**

ScRNA-seq sample sumaries before and after quality control filtering
d7m = day 7 minus dox, d7p = day 7 plus dox, d23m = day 23 minus dox, day 23 p = day 23 plus dox. (dox = doxycline induction of Hyp2 RNAi)

| sample | no. cell before QC | no. cell after QC | median UMI per cell | median features per cell | median % of transcrip | median % of transcripts encoding rRNA kDNA per cell |
|---|---|---|---|---|---|---|
| d7m | 18079 | 3826 | 1577 | 1389 | 1.174394426 | 1.382273421 |
| d7p | 10715 | 7721 | 1607 | 1363 | 0.924811301 | 2.485421341 |
| d23m | 9719 | 8435 | 1629 | 1333 | 1.542475956 | 2.742761405 |
| d23p | 8732 | 7590 | 1810 | 1428 | 1.630880926 | 2.057249839 |

QC threshold used for each sample

| sample | min. UMI | max. UMI | min. Feature | max. Feature | max. % kDNA | max. % rRNA |
|---|---|---|---|---|---|---|
| d7m | 1000 | 2100 | 1000 | 1720 | 2 | 3 |
| d7p | 900 | 2700 | 850 | 2000 | 1.8 | 5 |
| d23m | 500 | 3000 | 550 | 2100 | 2.9 | 5.5 |
| d23p | 500 | 3200 | 550 | 2200 | 2.9 | 4.5 |

Number of unique UMI (unique transcripts) per cell. Red dashed line indicates QC thresholds

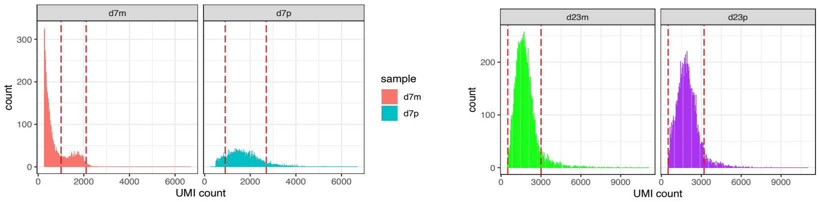


Number of unique features per cell. Red dashed line indicates QC thresholds

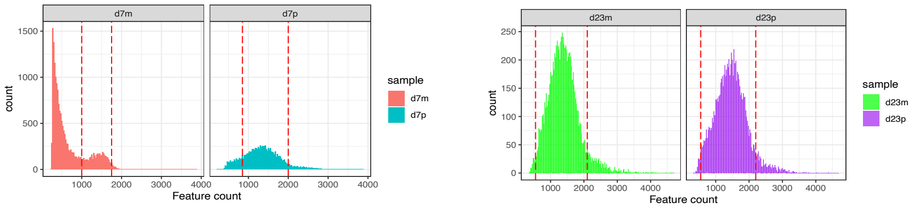


Percetage of transcripts per cell tgat are encoded on the kDNA maxicircle. Red dashed line indicates QC thresholds

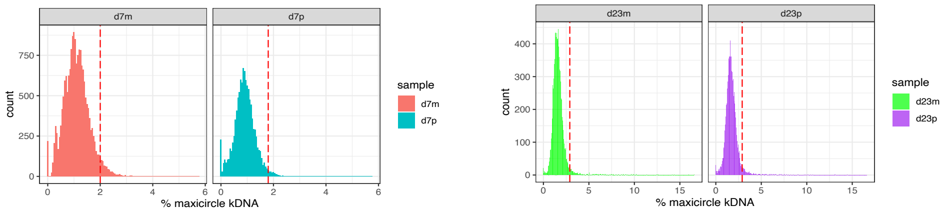


Percetage of transcripts per cell that encode ribosome RNA. Red dashed line indicates QC thresholds

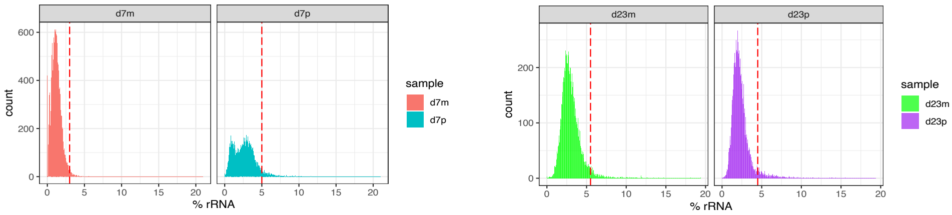


**Supplementary Figure 2**

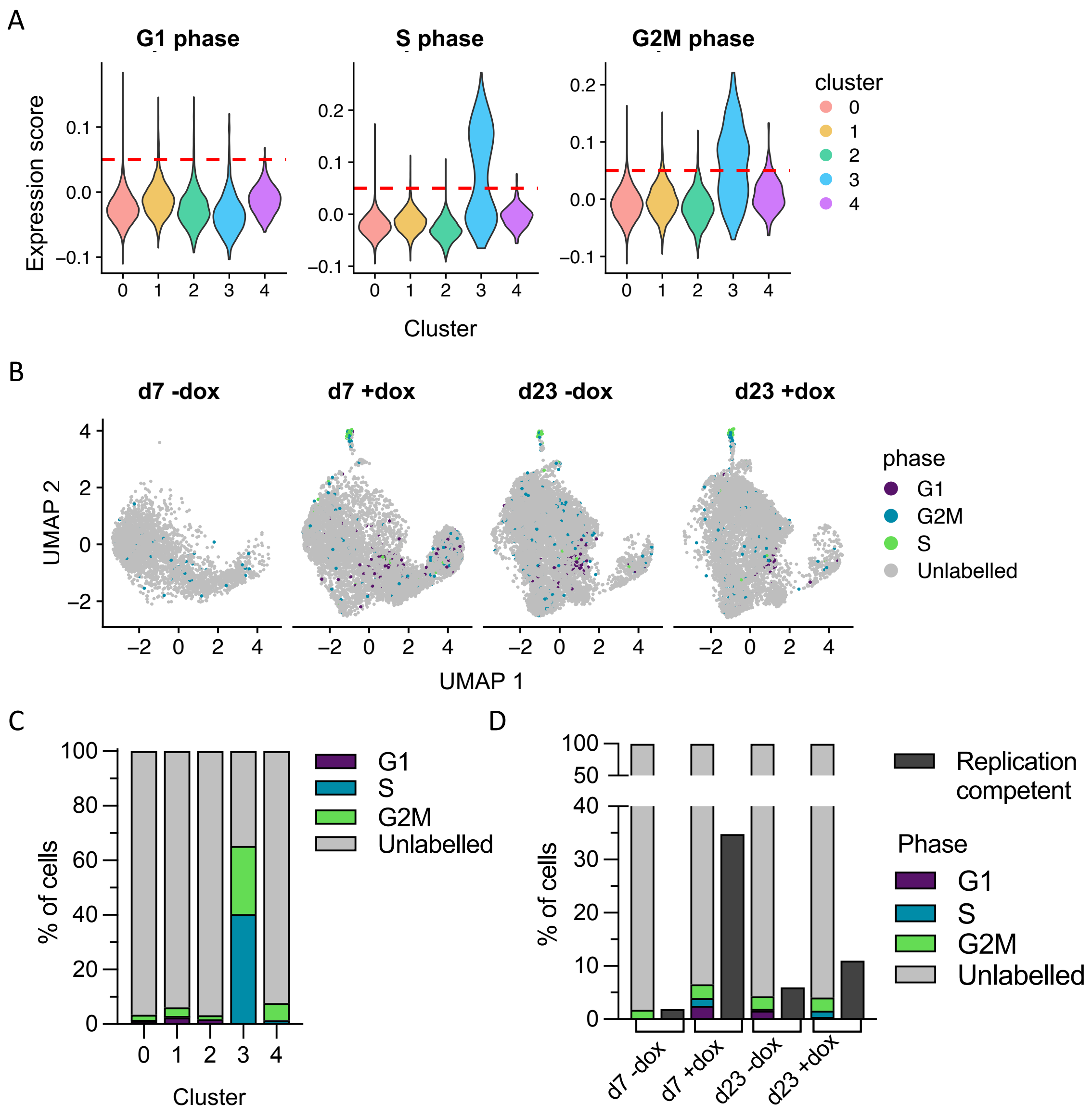


**Supplementary Figure 3**
